# SeqImprove: Machine Learning Assisted Creation of Machine Readable Sequence Information

**DOI:** 10.1101/2023.04.25.538300

**Authors:** Jeanet Mante, Zach Sents, Chris J. Myers

## Abstract

The progress and utility of synthetic biology is currently hindered by the lengthy process of studying literature and replicating poorly documented work. Reconstruction of crucial design information through post-hoc curation is highly noisy and error-prone. To combat this, author participation during the curation process is crucial. To encourage author participation without overburdening them, an ML-assisted curation tool called SeqImprove has been developed. Using named entity recognition, named entity normalization, and sequence matching, SeqImprove creates machine-readable sequence data and metadata annotations, which authors can then review and edit before submitting a final sequence file. SeqImprove makes it easier for authors to submit FAIR sequence data that is findable, accessible, interoperable, and reusable.

## Introduction

Synthetic biology has vast applications in various areas, including environmental, manufacturing, sensor development, defense, and medicine. However, the progress and utility of synthetic biology are currently hindered by the lengthy process of studying literature and replicating poorly documented work. The reuse of genetic components is currently low. ^1,2^ More complete data records makes the data more reusable and the database to which they are submitted more valuable.^3^

Much of the data that is submitted does not contain sufficient information for data reuse. The *Synthetic Biology Knowledge System* (SBKS) project attempted to address this via post-hoc curation.^4^ The project created an integrated knowledge system built using data generated with a post-hoc curation. The curation consisted of two parts: (1) a text mining that performs automatic annotation of the articles using *natural language processing* (NLP) to identify salient content such as key terms, relationships between terms, and main topics; and (2) a data mining pipeline that performs automatic annotation of the sequences extracted from the supplemental documents with the genetic parts used in them. The curation allows the linkage of knowledge, genetic parts, and the context in which they are used to the papers describing their usage. In order to process vast amounts of data, automated tools are employed to analyze unstructured text and identify relevant keywords, while attempting to derive their intended meaning from the surrounding context. This tests the limits of NLP methods, such as *named entity recognition* (NER) and *entity classification*. Additionally, it leaves ambiguous entities that only the original authors might disambiguate.^5^ For example, *S. aureus* may refer to *Scleropages aureus, Senecio aureus, Sericulus aureus, Somatogyrus aureus*, or *Staphylococcus aureus*. Furthermore, the SBKS project also extracted sequences provided as supplemental information in publications. However, these sequences, even when they are provided, are typically poorly annotated, incomplete, and provided in non-machine readable formats (e.g. PDFs). Taken together, the SBKS project demonstrated that reconstruction of important design information through post-hoc curation is extremely noisy and error prone.^6^

To address the issues raised by SBKS (NLP requiring manual refinement, ambiguity of language, and difficulty of sequence extraction), we suggest switching from post-hoc curation to *integrated curation*, i.e. asking authors to participate in curation before publication. The idea of author based curation (having the submitters curate their own data) is becoming increasingly popular.^7–9^ To enable author curation requires intuitive interfaces. The interfaces should prompt authors to ensure that all required metadata is present in predetermined locations and formats. Several different checklists for different experiment types have been suggested.^10–13^ However, these checklists do not have interfaces for authors and often do not specify a format, machine readable or otherwise, for required metadata. We developed the SeqImprove curation interface to provide an intuitive interface to allow author curation of machine readable sequence metadata. The interface allows authors to curate machine generated metadata and annotations and save the product of curation in a machine readable format. This paper presents the capabilities and underlying architecture of SeqImprove.

## Results

SeqImprove is designed to aid authors in creating machine readable sequence data with complete metadata. It consists of a user-interface that was built using modular code. It can be reused by others to work as the front-end for their curation software. Additionally, the back-end consists of a series of tools that automate NER, keyword grounding (*named entity normalization*) (NEN), sequence annotation, and protein prediction. The functions are accessed by users via the front end. The first step is sequence data input. As input, it takes in a sequence file in the *Synthetic Biology Open Language* (SBOL)^14^ format or a link to a sequence stored in SynBioHub.^15^ It then takes authors through 4 sections of metadata:

1. **Overview tab:** The first tab that the user encounters (Figure 1). It provides the description of the part (with all recognised terms hyperlinked out). Additionally, it allows users to select the role or function of the sequence via a drop down menu of *sequence ontology* (SO) terms.^16^ These terms include promoter, terminator, coding sequence, and more than 160 others from the sequence feature branch of SO. The next section allows users to link any target organisms, i.e. intended targets of sequence insertion. The machine readable formatting underlying target organisms is the NCBI Taxonomy.^17^ Species level organisms can be prioritised over strains and subspecies by selecting the “Prioritize parents in search” box. The final section of the tab is the references section. Any relevant papers or pre-prints may be linked using a DOI.
2. **Sequence tab:** This tab is the place where sequences are annotated with sub-components (Figure 2). The analyze sequence button can be used to generate suggestions of sub-components based on a SeqImprove’s library of frequently used components. The suggestions may be accepted by selecting the check box next to the sub-component’s name.
3. **Text tab:** This tab is where machine readable keywords are selected in the description field (Figure 3). The “Analyze Text” button uses machine learning to suggest keywords, group similar ones together, and suggest a machine readable ontology term for the keyword. Users can approve a keyword by selecting the checkbox next to it. They can also add terms that were missed to a group by highlighting the term and using the “Add to existing annoation” button that appears below the desicription box. New groups can be created using the “Create Text Annotation” button. The group label, identifier, and ontology term can be edited using the pencil button next to an annotation. Finally, the description can be edited using the pencil button at the top of the description box. This allows users to correct misspellings or add additional forgotten information.
4. **Proteins tab:** This tab is where proteins produced by the sequence are added to the metadata (Figure 4). There is a suggestion box on the right where proteins frequently associated with keywords or sequence annotations are provided. For example, *E. coli* in the description field leads to the suggestion of common *E. coli* proteins. The user can also add any further proteins from the UniProt database.^18–20^

After author review the file can be downloaded in the standard machine readable SBOL format. The file may then be shared with others, uploaded as part of a supplemental, or submited to SynBioHub.

**Figure 1:**
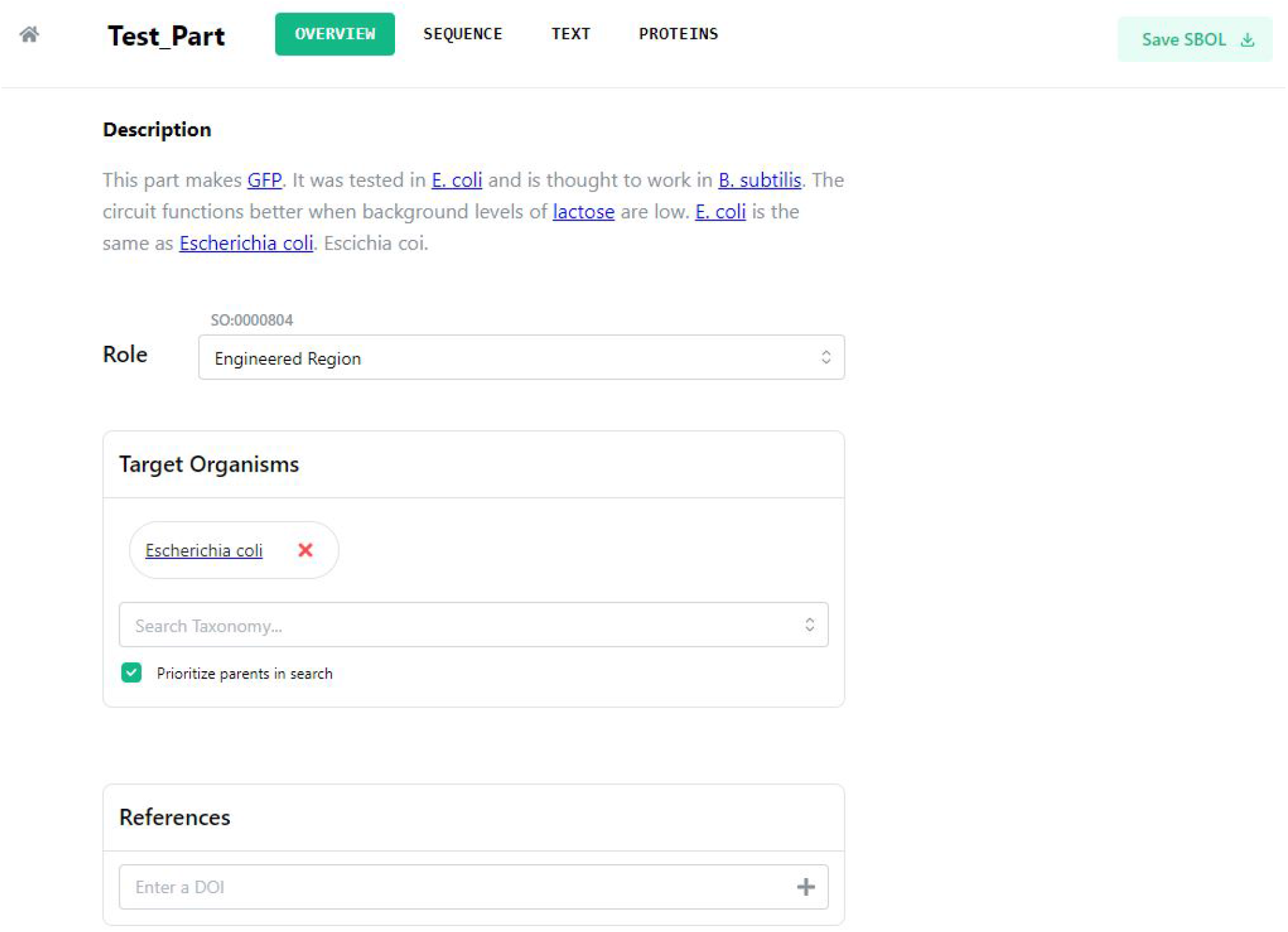
Overview Tab. This provides the user with a section where they can see the description and any recognised terms. They are also provided with the opportunity to define the role of the sequence they are submitting as well as the organism(s) it is designed for. Finally, any papers can be linked to the sequence via DOIs.

**Figure 2:**
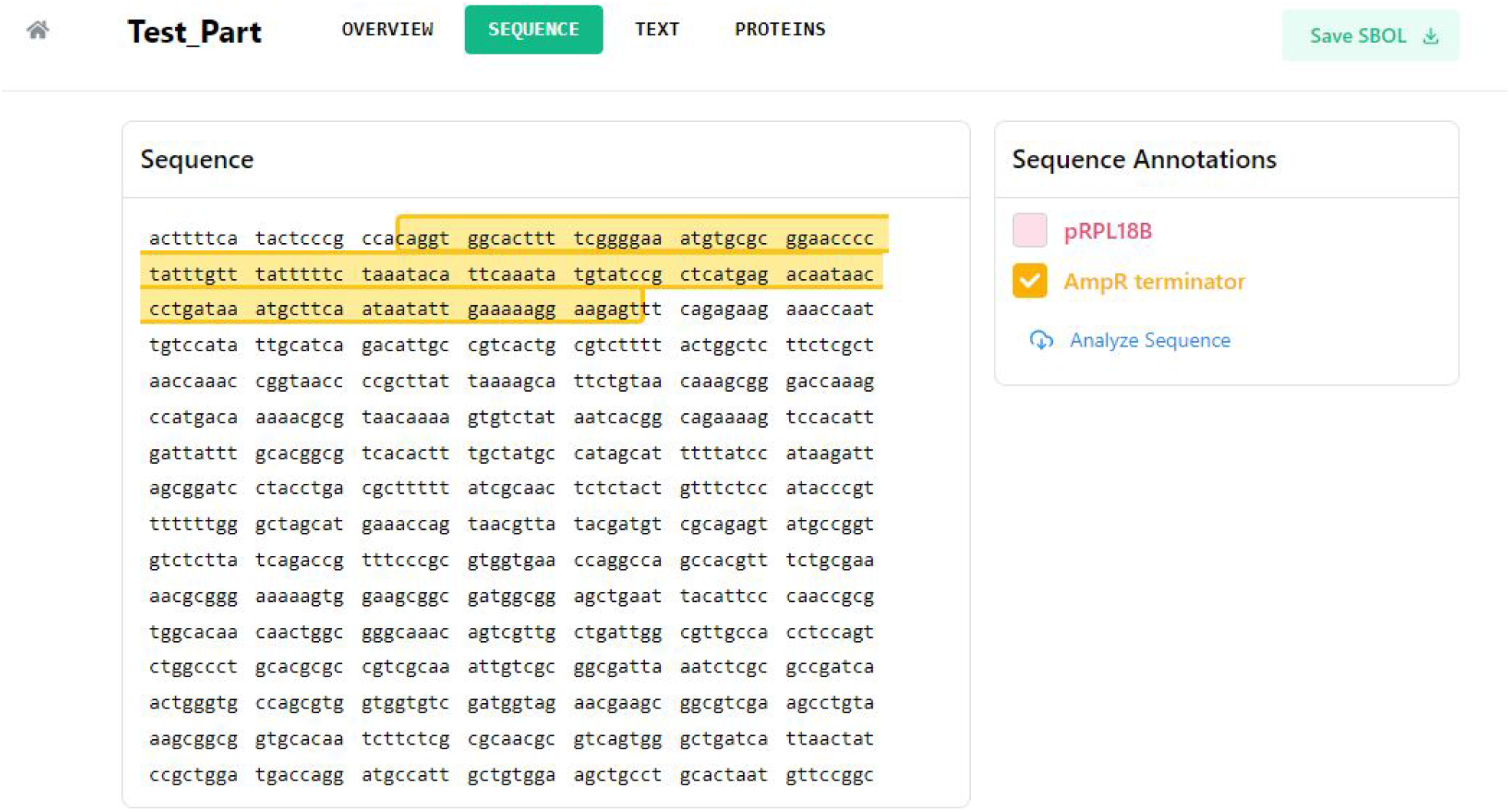
Sequence Tab. This provides sub-component suggestions for the sequence. The user may choose to approve the suggested sub-components by selecting the check box next to the sub-component names.

**Figure 3:**
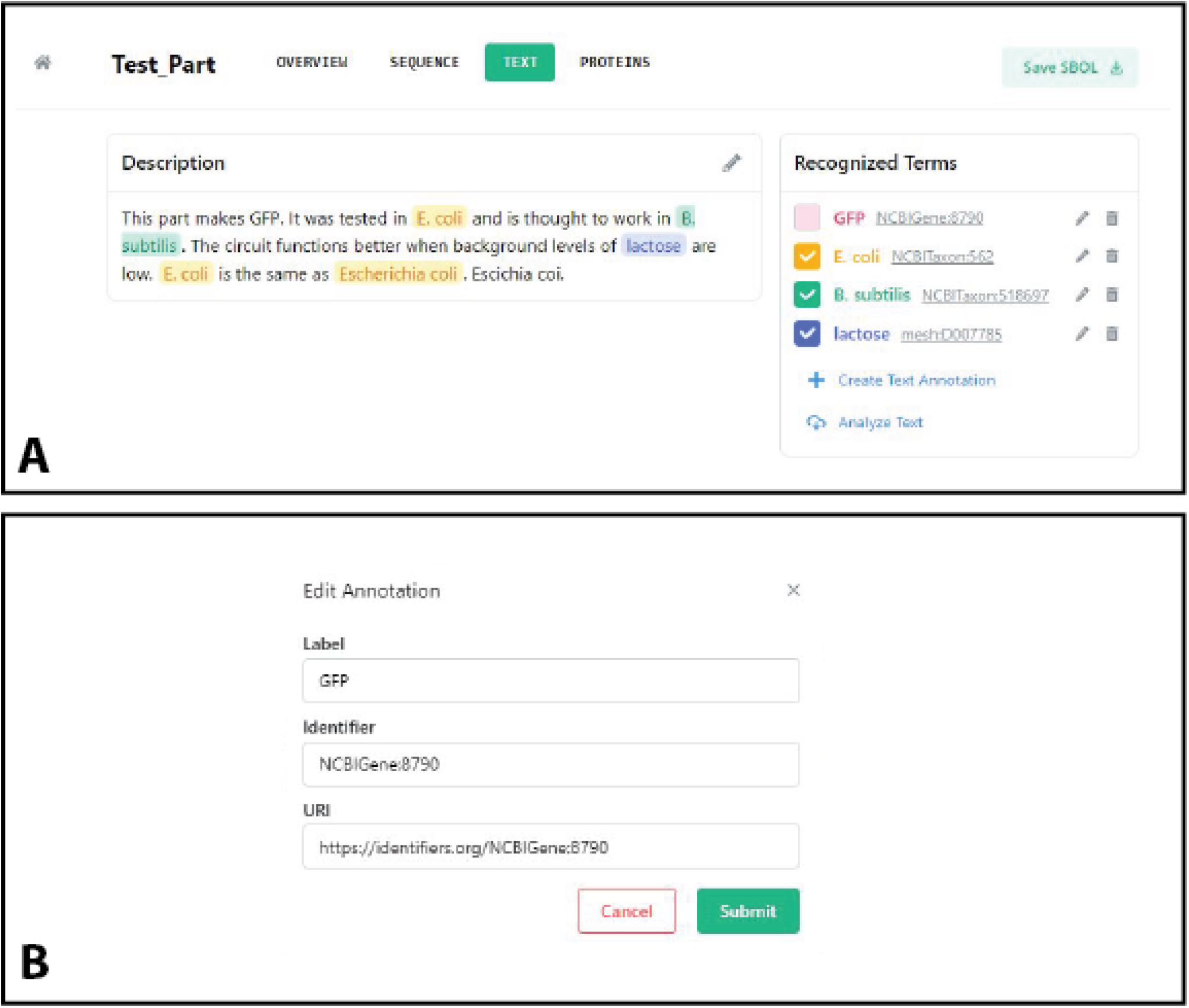
Text Tab. This tab is is used for adding recognised terms from the free text description. **A**: The desicription is shown with suggested key words. The description may be edited, to add more keywords or fix spelling mistakes. Additionally, suggested keywords may be accepted or rejected using the check boxes next to the term. Selection of text in the description box allows the addition of additional variations of a key word to existing recognised terms. **B**: Recognised terms can be edited. The human friendly name, identifier and full URI may be edited.

**Figure 4:**
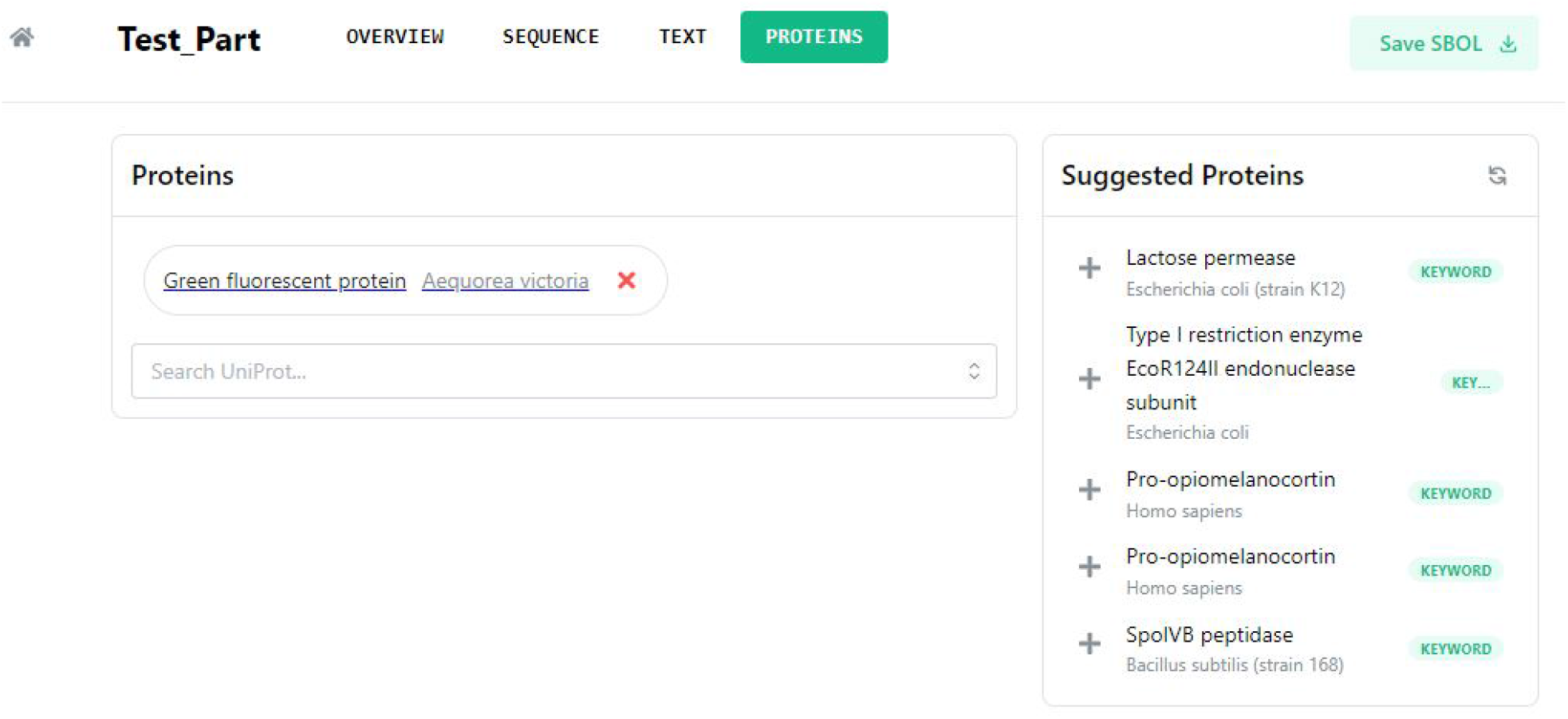
Proteins Tab. This tab allows linkage of proteins that will be produced by the sequence. The proteins link to UniProt IDs. There are suggestions on the right based on keywords or target organism matches. For example, as *E. coli* was recognised in the text common *E. coli*, proteins are suggested.

## Discussion

We have presented SeqImprove, a platform for machine-assisted author curation of genetic sequences. SeqImprove helps authors submit sequence data and associated metadata in machine readable formats. It prompts authors to consider metadata such as the: the role, target organism, reference papers, sub-sequences, protein production, and keywords in their free text description. It makes the information machine readable by using existing ontologies to structure the metadata. Authors are helped by suggestions of keywords, proteins, and sequence annotations. They can review and edit the suggestions in a user friendly interface. The interface was also designed to be modular so it could be reused for similar editing and review in other curation contexts.

While SeqImprove offers many benefits there are still limitations to the system for future work to address. The first limitation is author participation. SeqImprove only works if authors use it. Despite the major benefits offered by SeqImprove, it can be difficult to incentivize researchers to participate. The benefit for a researcher is the future reduction of time to find sequences that others have submitted and, hopefully, more citations to their own easier to find sequences. Thus, the results of additional effort in curation are not immediate and a little indirect. This is particularly the case initially, as there will be little well curated output data to be utilized. This reduces the incentives for researchers to adopt the system, and without adoption, the available data remains scarce. Breaking the initial consensus threshold will be difficult, and would be aided by journal incentives. We are working towards agreements with journals to add requirements for sequences to be submitted via curation systems such as SeqImprove.

A second major limitation is the limited abilities to leverage better curated files via enhanced search algorithms. We are working towards a database that utilizes the curated sequence files to make answering research questions easier. Curation underpins natural language queries with query refinement based on the metadata, e.g. suggesting different taxonomic terms for a query with a species name if no results or too many results are found. Providing databases with curation enabled search will also increase the incentives for SeqImprove use.

A third suggested improvement is research on the metadata required for component reuse. SeqImprove currently collects information based on a qualitative survey of article methods sections and sequence databases. Research to create a minimum information standard and characterisation protocols for synthetic biology sequences is still required. Such a standard would drive the information collected by SeqImprove.

Further improvements are less conceptual. The editor should be expanded to allow new file creation as well as editing and adding metadata to current files. It should also be expanded to support other input formats, such as GenBank files with the help of the GenBank to SBOL Converter.^21^ Additionally, to make submission and curation workflows more streamlined, the editor will be linked to SynBioHub via plugins. ^22^ This will allow users to access the editor both independently and as part of SynBioHub workflows. Furthermore, the sequence annotation tab will be expanded. Additional features will include authors being able to add their own sequence annotations, similar to the keyword additions, and expanding the methods by which sequence annotations are suggested, e.g. via BLAST matches. Finally, all of the machine learning models will be improved over time. This can be done by using sequences curated by SeqImprove users to expand the training sets of the models.

## Methods

SeqImprove is an application for curating genetic designs encoded in SBOL. It can be run standalone or as a SynBioHub curation plugin. ^6,22^ It is meant to help users easily add metadata to their genetic designs by providing recommendations and a simple interface with which to do so. SeqImprove consists of two applications and a package. The two applications are a React front-end and a Dockerized Express/Node.js API that functions as the back-end. The package is called text-ranger and was developed to make working with text ranges and replacements easier, as this is key functionality for creating and displaying text annotations for genetic designs.

The backend has two main machine aided curation functions:

1. **Annotate Sequence**: This is the method used to suggest sequence annotations. It is based on SYNBICT.^23^ It uses the feature libraries found in the Github Repository. These include libraries from parts-rich papers.^24–31^
2. **Annotate Text**: This method is used to suggest keyword annotations. It uses BERN2 for NER and NEN (i.e. grounding).^32^ Additional fuzzy matches are carried out to catch potential misspellings using the fast-fuzzy NPM package.

Text-ranger is an internal Node.js package written to aid in the modeling of text annotations. It provides an interface for creating text replacements based on a start and end position. It compiles the replaced text on demand and adjusts replacement positions if the underlying text is edited. This package is used as part of the front-end for text annotation.

## Supporting information

Demo Video

## Acknowledgement

JM, and CM are supported by the National Science Foundation under Grant No. 1939892. JM and ZS are supported by the National Science Foundation under Grant No. 2231864. This document does not contain technology or technical data controlled under either U.S. International Traffic in Arms Regulation or U.S. Export Administration Regulations. Any opinions, findings, and conclusions or recommendations expressed in this material are those of the author(s) and do not necessarily reflect the views of the funding agencies. All authors contributed to the writing of this manuscript.

## Author Contributions

All authors contributed to the writing of this manuscript. JM designed the high level architecture for SeqImprove. ZS implemented the design and wrote the code. CM provided guidance throughout. All authors contributed to the writing of this manuscript.

## Supporting Information Available

- GitHub: https://github.com/MyersResearchGroup/SeqImprove

## Suplemental Files

- Demo Video.vid

## TOC Graphic

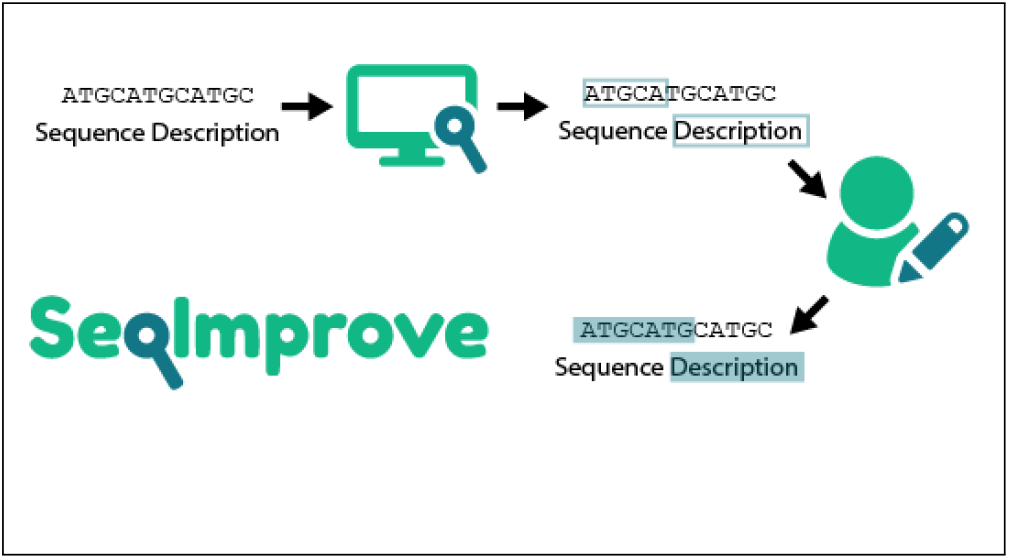

